# An agent-based 3D model of non-genetic adaptation in cancer tissues under electrical, mechanical, and hypoxic stress

**DOI:** 10.64898/2026.08.31.748266

**Authors:** João Ferreira Gil, Nathalia Pinheiro, Saverio Gentile, Gil Gonçalves, Rosalia Moreddu

## Abstract

Non-genetic adaptation enables cancer cells to alter their phenotype under stress without requiring new mutations. However, the mechanisms by which electrical, mechanical, and hypoxic cues combine to shape this process in 3D tissues remain poorly understood. This work presents an agent-based tumor model that integrates vascular oxygen supply, a globally imposed electric field, mechanically mediated crowding and compression cues, phenotype transitions, cell growth, mitosis, death, and inheritance of adaptive memory across division. The simulated tumors exhibit a three-stage trajectory consisting of necrosis onset, transient collapse of live mass, and partial regrowth accompanied by progressive accumulation of adapted cells. Continuous electrical stimulation produces a dose-dependent reduction in live mass while markedly increasing the adapted fraction, with comparatively limited changes in final necrotic burden. This response is strongly conditioned by mechanics and reshapes (and is reshaped by) adaptive capacity. Pulsed stimulation further shows that, in the model, electric field amplitude and temporal schedule jointly determine memory phenomena, phenotypic diversification, and growth recovery. These results show that coupling local oxygen availability, mechanical constraints, electrical forcing, and history-dependent phenotype transitions can generate distinct tissue-level patterns of phenotypic heterogeneity. Both stimulus magnitude and temporal protocol influenced the resulting population structure, suggesting that the history of physical stress may be an important determinant of adaptive dynamics in spatially organized tumor models.

## 1. Introduction

Cancer progression emerges from a continuous interaction between malignant cells and the microenvironments they create and inhabit^1, 2^. Cancer heterogeneity is central to how tumors grow, invade, metastasize, and respond to therapy^3, 4^. Clinical failure often reflects the capacity of cancer cell populations to reorganize under stress^5, 6^, because malignant cells can alter their behavior without acquiring new genetic lesions^5, 7^. To understand tumor progression, cancer should also be viewed as a spatially organized adaptive system in which local environmental conditions help determine cell fate, survival, and population structure over time. At the heart of this adaptive capacity is non-genetic adaptation, also called phenotypic plasticity^8^. Tumor cells can shift between plastic, quiescent, stressed, migratory, and stem-like states in response to environmental signals, and these transitions can occur on timescales much faster than clonal evolution^9^. Some shifts are reversible, whereas others persist long enough to create functionally stable subpopulations, suggesting that short-term adaptation can lead to long-term selection^10, 11^. Plasticity therefore links cell-scale physiology to population-scale heterogeneity. A deeper understanding of these dynamics could reveal complementary disease mechanisms, help identify new therapeutic targets, and inform the design of bioinspired artificial systems^12, 13^. Among the microenvironmental drivers of tumor behavior, oxygen availability is one of the most pervasive and best established^14^. Tumor vasculature is typically disordered, tortuous, and inefficient, producing uneven delivery of oxygen and nutrients. These conditions reshape metabolism, alter cell-cycle progression, promote angiogenic signaling, influence genomic stress responses, and, when severe, lead to necrosis^14^. Oxygen is also a powerful organizer of space: cells near supply sources may remain plastic, whereas deeper regions become stressed or dormant^15^. Because such patterns emerge through diffusion, consumption, and geometry,^14^ homogeneous descriptions of oxygen can miss the internal structure that drives cell behavior in 3D tissues.

Mechanical forces are equally important and are harder to quantify experimentally^16^. Optomechanical methods used as physical strain sensors^17^ have been refined to interface contracting cells to record biomechanical activity^18^. However, such approaches are impracticable in mostly stationary cells where mechanical interactions happen within cells across 3D constructs, exerting minimal forces of harder access. As tumors expand within confined tissue, they generate compressive stress, remodel the extracellular matrix, and alter cell-cell and cell-matrix force transmission^19^. Changes in tumor stiffness, confinement, and crowding can regulate proliferation, apoptosis, migration, polarity, and invasive behavior through mechanosensitive pathways that act from the membrane to the cytoskeleton and nucleus^20^. Mechanical stress can also reshape the biochemical landscape by compressing vessels, thereby intensifying oxygen limitation^21^. Conversely, altered oxygenation and metabolism can modify matrix remodeling and tissue architecture^22^. Electrical phenomena, such as variations in membrane potential, ionic fluxes, and endogenous electric fields, represent a third dimension associated with cellular adaptation^23-25^. Although the literature on tumor bioelectricity is still less mature than work on cancer hypoxia, mechanics, or electrophysiology work in excitable tissues^26-28^, it is now established that dysregulated ion channel expression in cancer cells is associated with malignant behavior, metastatic potential, collective coordination, and treatment response^24, 25^. These classes of cues (oxygenation, mechanics, and electrical state) operate jointly, and often non-linearly. Experimental approaches have revealed many of these principles, but directly observing their interaction in living tumor tissue remains difficult. Therefore, computational models have emerged as powerful complements^29, 30^. Continuum models efficiently capture diffusion, transport, and large-scale tissue mechanics, but they typically average away the discrete competition and cell-state variability that drive adaptation at the microscale^31^. Intracellular network models can describe signaling logic in detail, but they are rarely embedded in evolving spatial environments^32^. Agent-based approaches are a useful middle ground because they represent individual cells as units that sense local conditions, change their phenotype, divide, die, and compete for space^33^. Despite these advantages, many agent-based studies examine oxygen alone, mechanics alone, or a single signaling cue superimposed on generic growth dynamics^34^. Others represent phenotypic variation only as fixed cell types, without allowing history-dependent switching or inheritance of adaptive state variables across division^35^.

The present study reports a 3D agent-based tumor model to examine how vascular oxygen supply, electrical signaling, and mechanical interactions jointly influence tumor growth and non-genetic phenotypic diversification (**Figure 1**). The model integrates tissue initialization, a spatial oxygen field linked to vascular input, an electrical field module, a mechanical cue module, phenotype update rules, mitosis, cell death, and inheritance of adaptive state variables. This architecture enables to ask whether coupled microenvironmental pressures are sufficient to generate emergent behaviors that are not predictable from isolated cues alone, including altered growth kinetics, spatial reorganization, shifts in phenotype composition, and persistence of adapted states. By linking local microenvironmental cues to cell-level behavior in a 3D setting, this study aims to investigate how adaptive heterogeneity can arise as a system property and provide a versatile tool for further multiscale investigations in 3D tissues. Within the model, reversible entry into the P state represents plasticity, whereas transition from P to A represents stable adaptation with inherited stress-memory traces across division.

**Figure 1.**
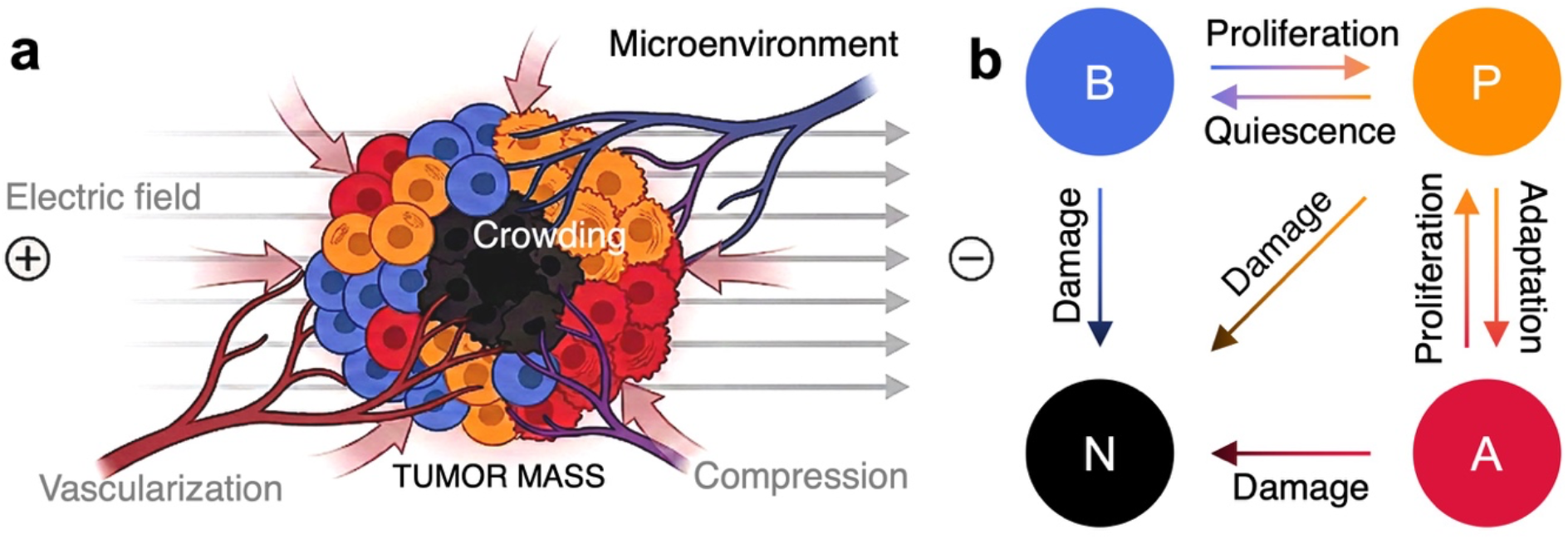
Model architecture coupling electric, vascular, and mechanical cues to tumor-state transitions. **(a)** Schematic of a vascularized tumor mass embedded in a heterogeneous microenvironment and exposed to a left-to-right electric field. The framework integrates vascular support with spatially resolved crowding and compression within the tumor. **(b)** State-transition map for baseline (B), plastic (P), adapted (A), and necrotic (N) cells. Reversible transitions encode phenotypic plasticity and adaptation, whereas damage-driven transitions lead to cell necrosis.

## 2. Results

### 2.1. Reference full-model dynamics show early collapse followed by spatial reorganization and adaptive regrowth

Simulations of the reference full-model condition (continuous E3 with mechanics and adaptation enabled, see *Methods*) resolved a three-stage response (**Figure 2**). Throughout this manuscript, adaptation refers to the model-defined A state. The phenotype renders in Figure 2a display that the early state retained a largely baseline architecture, with plastic cells concentrated near the outer surface and only a small necrotic core emerging centrally. The oxygen maps already showed a sharp supply gradient at this point, with the brightest signal confined to the exposed outer rim and progressive depletion toward the center (Figure 2b), indicating that the interior became resource-limited before structural failure was complete. In parallel, the voltage field imposed a stable left-to-right polarity across the aggregate throughout the time course (Figure 2c).

**Figure 2.**
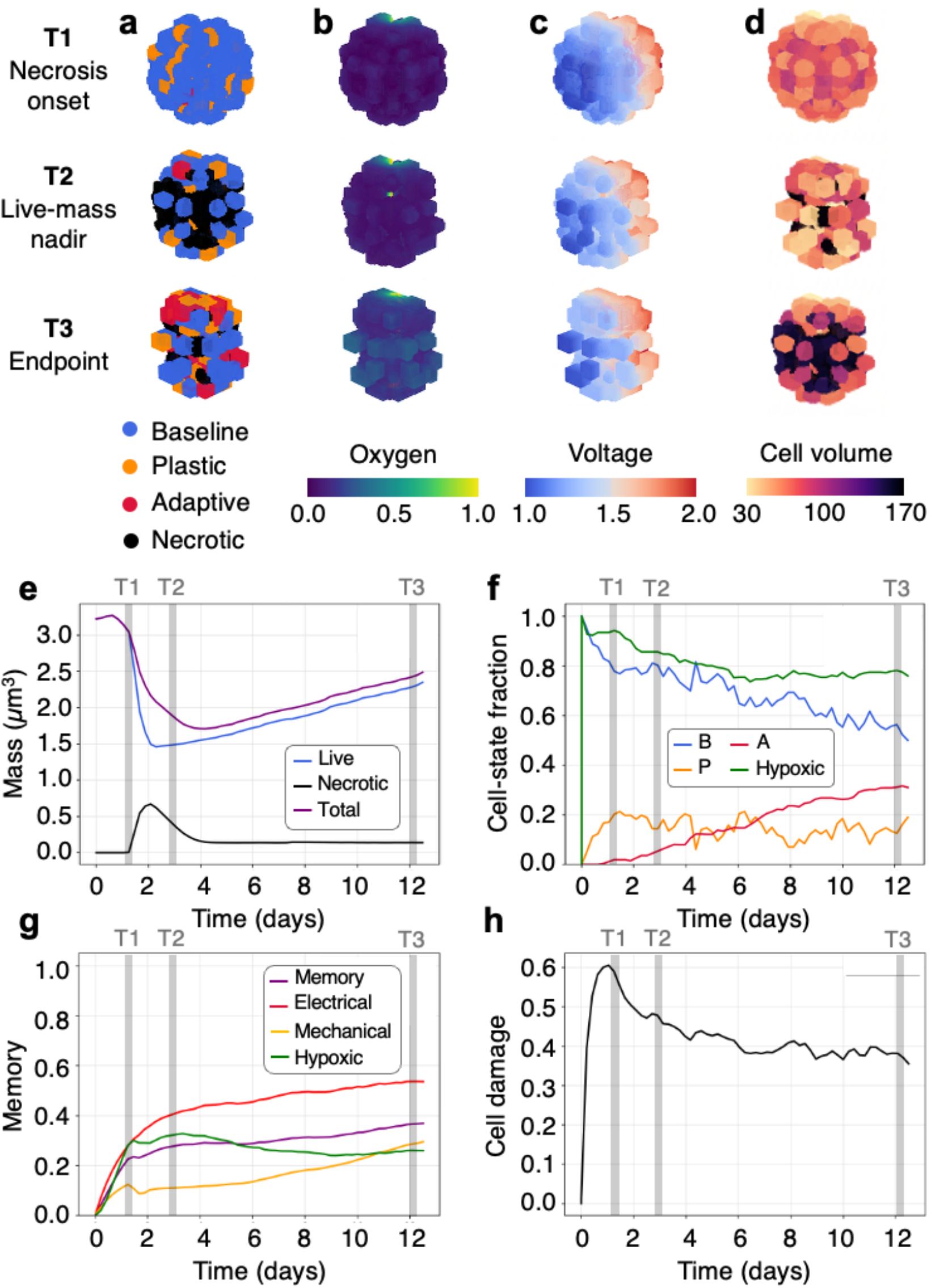
Reference full-model dynamics. **(a-d)** Renders at landmark time points T1 (necrosis onset), T2 (live-mass nadir), and T3 (endpoint), showing cell-state composition (**a**), oxygen (**b**), voltage (**c**), and cell volume (**d**). **(e-h)** Corresponding time courses of live, necrotic, and total mass (**e**), cell-state fractions and hypoxic fractions (**f**), total and component memories (**g**), and mean cell damage (**h**). Gray bands mark T1–T3.

By T2, these cues had induced morphological specialization: necrotic channels expanded through the core, baseline cells became segmented, and adaptive and plastic populations accumulated preferentially along the periphery where oxygen access was greatest. In the current implementation electrical stress magnitude is spatially uniform during on phases (Figure 2a-c). By T3, the spheroid had stabilized in a remodeled configuration characterized by a persistent necrotic scaffold, a broader adaptive shell, and a more heterogeneous cell-volume distribution at the outer boundary (Figure 2d). The overall response shows a spatially structured transition toward a new steady architecture.

The temporal trajectories quantify this restructuring and show that the morphological phases corresponded to a sequence of collapse and rebound. Live mass fell from approximately 3.1 to 1.5 before recovering to about 2.35 by the endpoint, while total mass followed a similar path from roughly 3.2 to 1.75 and then rose to about 2.45 (Figure 2e). Necrotic mass increased from zero to a peak near 0.65 around the early-middle interval and then declined toward 0.1, indicating that the deepest injury phase preceded the regrowth phase rather than persisting throughout it. This pattern shows that the tissue did not recover by tolerating an ever-larger necrotic burden. Instead, it partially cleared or compacted the damaged compartment while rebuilding the viable rim. The state fractions support this interpretation. Baseline occupancy dropped from about 0.90 to 0.52, plastic cells rose transiently to roughly 0.20 and then stabilized near 0.15, and adaptive cells accumulated steadily to about 0.31 by the endpoint (Figure 2f). Simultaneously, the hypoxic exposure fraction decreased from nearly 1.0 to approximately 0.76, indicating that survivors progressively redistributed toward more favorable microenvironments. Endpoint recovery reflected compositional rewriting of the spheroid.

The memory variables (see *Methods*) show that this rewriting was history-dependent and mechanistically layered. Electrical memory rose most strongly, reaching approximately 0.54 by the endpoint, whereas mechanical memory accumulated more gradually and plateaued near 0.27 Hypoxic memory increased early, peaked near 0.32, and then settled around 0.25, preserving a durable imprint of the initial oxygen crisis even after the most acute phase had passed. Integrated memory rose more slowly to about 0.36, consistent with it acting as a cumulative summary of the other channels (Figure 2g). Cell damage showed the earliest and steepest excursion, climbing to roughly 0.60 and then declining to around 0.37 by the endpoint (Figure 2h). The ordering of these curves shows that damage is registered first, electrical memory became the dominant persistent signal, mechanical memory matured during structural remodeling, and integrated memory consolidated later. The significance of this pattern is that the final phenotype encoded the sequence of stresses experienced during collapse and regrowth. Baseline recovery therefore emerged as a memory-bearing process in which electrical, mechanical, and hypoxic histories remained visible in the endpoint state.

### 2.2. Electrical response depends on mechanics and adaptation

To determine how mechanical signaling and adaptive competence shape field response, we compared the four endpoint conditions M0/A0, M0/A1, M1/A0, and M1/A1, corresponding to on/off states of mechanics and adaptation in the simulation (**Figure 3**). At E0 with electric field module off, live mass separated into two regimes: the mechanics-off states clustered at 26.79×10^6^ and 26.62×10^6^, whereas the mechanics-on states were lower at 24.71×10^6^ and 24.37×10^6^ (Figure 3a). This initial split indicates that mechanics alone constrained bulk expansion. Under mid-range electric field, the ranking changed subtly but the same broad structure remained. M0/A1 preserved the highest live mass at 26.43×10^6^, followed by M0/A0 at 25.92×10^6^, whereas M1/A0 and M1/A1 ended lower at 23.18×10^6^ and 22.85×10^6^, respectively. The E0-E3 live-mass shifts were therefore modest in M0/A1, larger in M0/A0, and strongest in both mechanics-on states, showing that mechanical coupling increased sensitivity of viable bulk to sustained electric field exposure. Importantly, the necrotic mass behaved differently.

**Figure 3.**
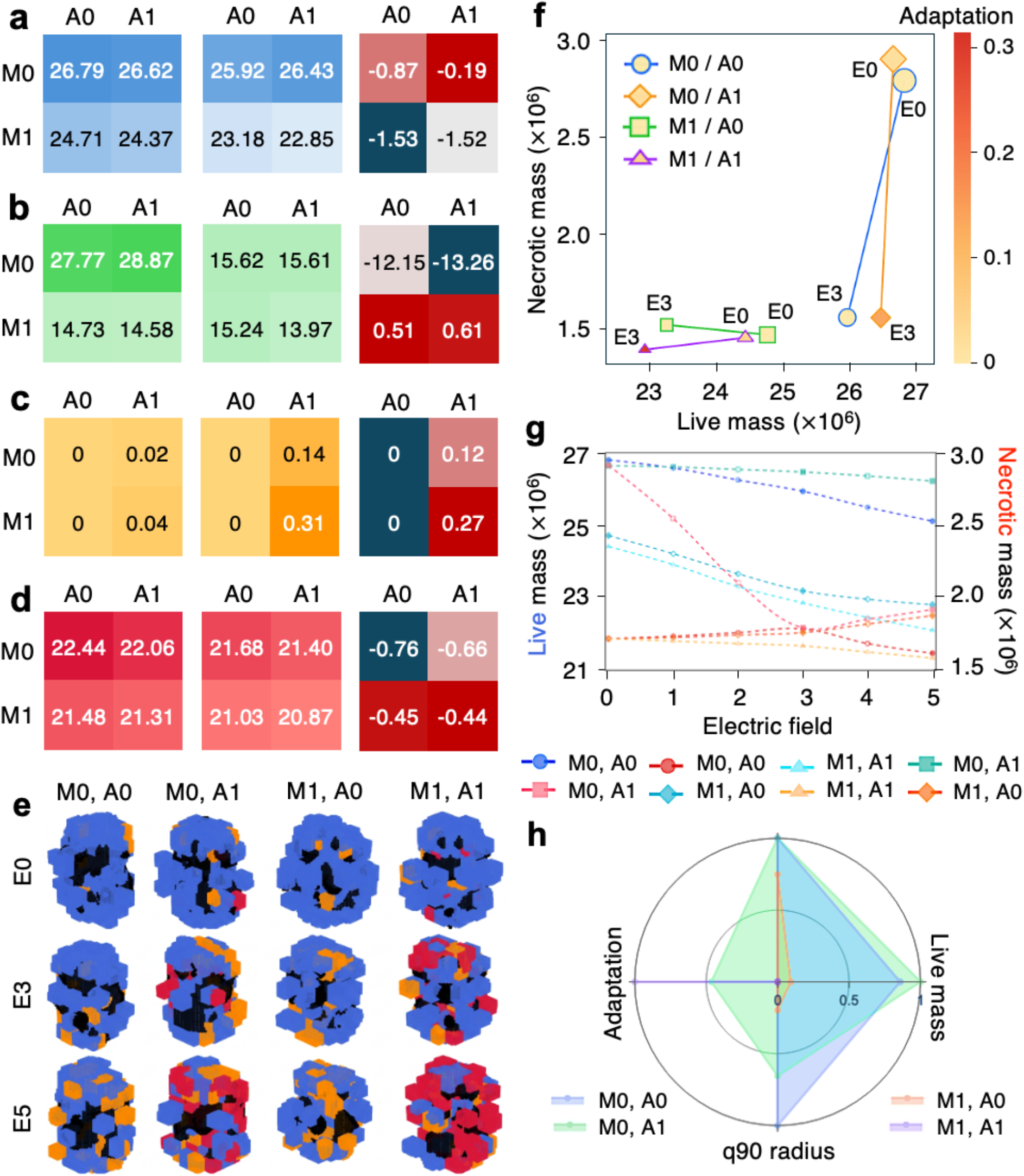
Response to electric fields depends on mechanics and adaptive capacity. **(a–d)** Heat maps summarizing endpoint live mass (**a**), necrotic mass (**b**), adapted fraction (**c**), and q90 radius (**d**; 90th percentile of live-cell radial distances) for the four combinations of mechanics off/on (M0/M1) and adaptation off/on (A0/A1). Left and middle matrices show E0 and E3, respectively, and right matrices show Δ(E3−E0). In a-d, values are scaled by a factor of 10^6^. **(e)** Endpoint 3D cell-state renders for all M/A conditions at E0, E3, and E5. **(f)** Live-mass versus necrotic-mass trade-off colored by adapted fraction; trajectories connect increasing electric field strengths within each condition. **(g)** Endpoint live mass (left axis) and necrotic mass (right axis) across electric-field amplitude. **(h)** Radar summary of normalized live mass, q90 radius, and adaptation across conditions.

At E0, M0/A0 and M0/A1 occupied a high-necrosis branch at 2.78×10^6^ and 2.89×10^6^, while M1/A0 and M1/A1 were much lower near 1.47×10^6^ and 1.46×10^6^ (Figure 3b). By E3, these values converged into a narrower 1.40–1.56×10^6^ band. This indicates that field exposure compressed the differences in necrotic burden while preserving differences in live mass. The remaining endpoint metrics clarify how the same four conditions resulted in diverse mass states. As expected, the A0 conditions remained at zero adapted fraction across both fields (Figure 3c). In contrast, M0/A1 increased from 0.02 at E0 to 0.14 at E3, whereas M1/A1 increased from 0.04 to 0.31, making M1/A1 the most adaptation-rich branch by a substantial margin. The q90 radius decreased in every condition, from 22.4 to 21.7 in M0/A0, 22.1 to 21.4 in M0/A1, 21.5 to 21.0 in M1/A0, and 21.3 to 20.9 in M1/A1 (Figure 3d) indicating that electric stimulation and mechanics both contributed to compaction. Representative 3D phenotype renders support this interpretation visually: E0 conditions appeared more baseline-dominant with compact central necrotic regions, whereas E3 conditions showed stronger peripheral plastic and adaptive enrichment, particularly when adaptation was enabled (Figure 3e,f). The multivariate summaries then sharpen the distinction among regimes. In the live vs. necrotic plane, M0/A1 combined the best preservation of live mass with marked necrotic reduction, while M1/A1 paired the smallest overall footprint with the largest adaptive fraction (Figure 3g). The radar comparison confirms the same partitioning, identifying M0/A1 as the mass-preserving solution and M1/A1 as the adaptation-maximizing solution (Figure 3h). Therefore, in this model, mechanics and adaptation together redirect the spheroid toward different response strategies.

### 2.3. Higher electric field strength drives transition toward adaptive states

An electric field-amplitude sweep within the enabled M1/A1 condition shows a dose-response structure that was largely monotonic (**Figure 4**). Live mass declined almost linearly from about 24.8×10^6^ at E0 to 21.7×10^6^ at E5, corresponding to an average loss of roughly 0.62×10^6^ per field increment (Figure 4a). By contrast, necrotic mass remained confined to a narrow range of approximately 1.38–1.57×10^6^ across the entire sweep (Figure 4b). This difference is biologically meaningful because it shows that stronger fields progressively reduced the viable, growth-supporting compartment while keeping total necrotic burden relatively constrained. Adapted fraction increased from about 0.04 at E0 to roughly 0.54 at E5, with a steeper rise from E3 onward (Figure 4c). At the same time, the q90 radius decreased from about 21.35 to 20.78 (Figure 4d), indicating that higher fields yielded smaller and denser spheroids. E3 occupied a particularly informative midpoint in this progression: by that amplitude, live mass had already fallen to about 22.9×10^6^, necrotic mass remained near 1.40×10^6^, adapted fraction had risen to around 0.31, and q90 radius had contracted to about 20.9 (Figure 4a–d).

**Figure 4.**
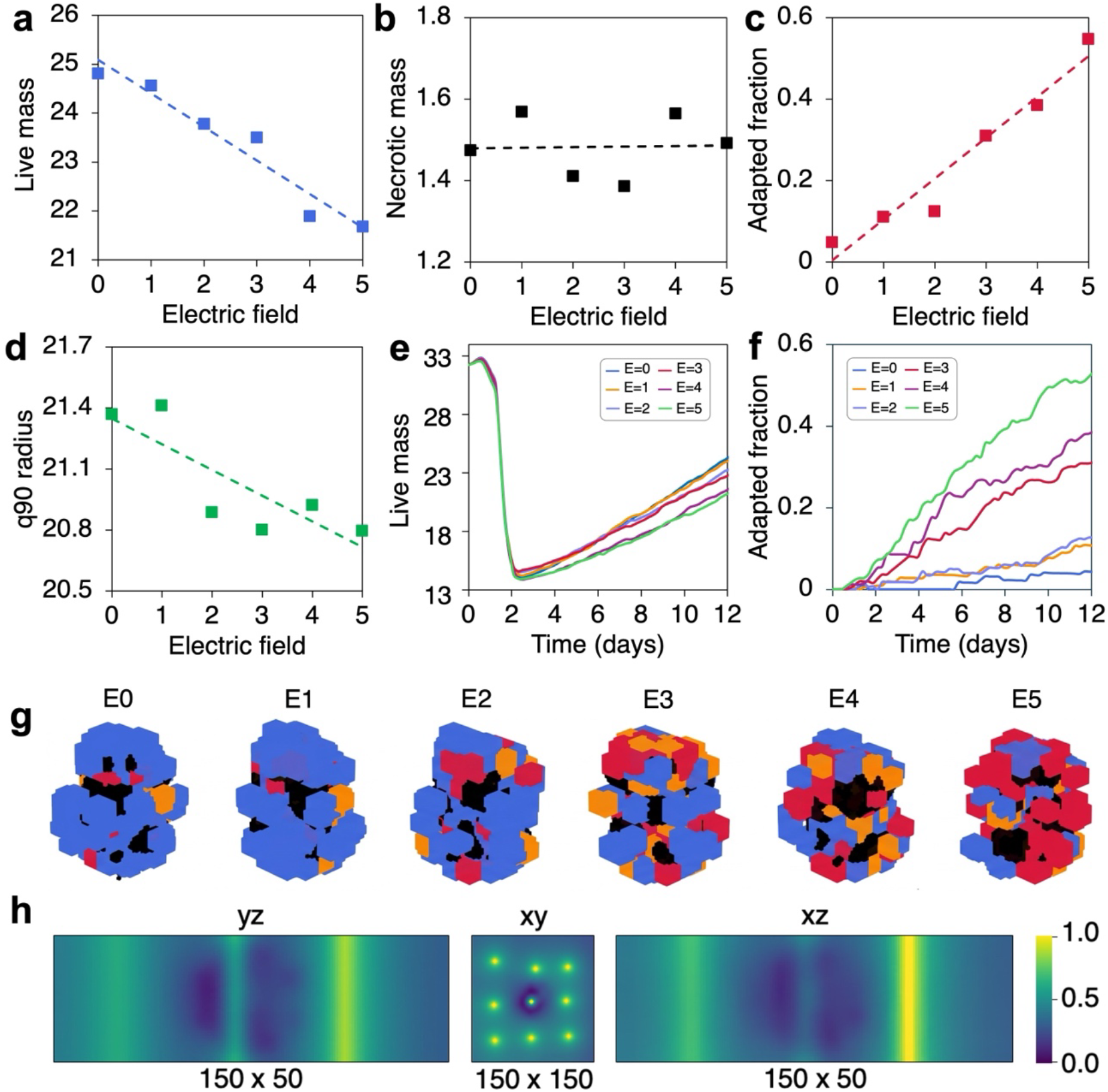
Continuous electric field dose response in the full model shows that increasing electric field reduces tumor mass while enriching adapted cells. **(a-d)** Endpoint live mass (**a**), necrotic mass (**b**), adapted fraction (**c**), and q90 radius (**d**) as functions of electric-field amplitude; dashed lines indicate linear fits. **(e, f)** Time-resolved live mass (**e**) and adapted fraction (**f**) for the E0–E5 series. **(g)** Endpoint 3D cell-state renders across the field range. **(h)** Orthogonal yz, xy, and xz views of the fixed vascular/oxygen-source geometry used to define the microenvironment.

The time-resolved and spatial panels explain how these endpoints emerged. All live-mass trajectories shared the same early collapse, dropping from roughly 33 to 14 by around day 2, after which they diverged smoothly during regrowth (Figure 4e). E0 and E1 formed the upper recovery branch, E2 and E3 occupied an intermediate branch, and E4 and E5 stabilized lowest, producing an endpoint spread of about 2.5 units. Adapted-fraction trajectories separated even more sharply: E0–E2 rose slowly and finished near 0.03–0.12, whereas E3, E4, and E5 ended near 0.31, 0.38, and 0.52, respectively (Figure 4f). This finding identifies the transition between E2 and E3 as that where the largest increase in adapted fraction occurred. The 3D renders translate this threshold into morphology. Low-field spheroids remained dominated by baseline cells, intermediate fields introduced clearly visible adaptive sectors at the periphery, and high fields produced extensive adaptive shells surrounding enduring necrotic channels (Figure 4g). The oxygen cross-sections shows that all amplitudes retained bright peripheral supply and dark central depletion, but the adaptive outer layer became increasingly prominent around these chronic oxygen gradients as field strength increased (Figure 4h). The result is that stronger fields reallocate the tissue into a more adaptive and strongly surface-organized state.

### 2.4. Pulsed electrical stimulation produces schedule-dependent and context-dependent adaptation outcomes

Because continuous amplitude is only one dimension of control, the next step is meant to study how pulsed schedules reshape endpoint composition (**Figure 5**). The pulse designs are summarized in Figure 5a, including the reference 240/180 period/off pattern and the four variants P1 (240/120), P2 (240/60), P3 (120/60), and P4 (60/30). Under the reference protocol, endpoint composition across the four condition classes and three field amplitudes spanned wide but structured ranges: baseline counts varied from 71 to 167, plastic counts from 7 to 34, adaptive counts from 0 to 52, and necrotic counts from 149 to 178.

**Figure 5.**
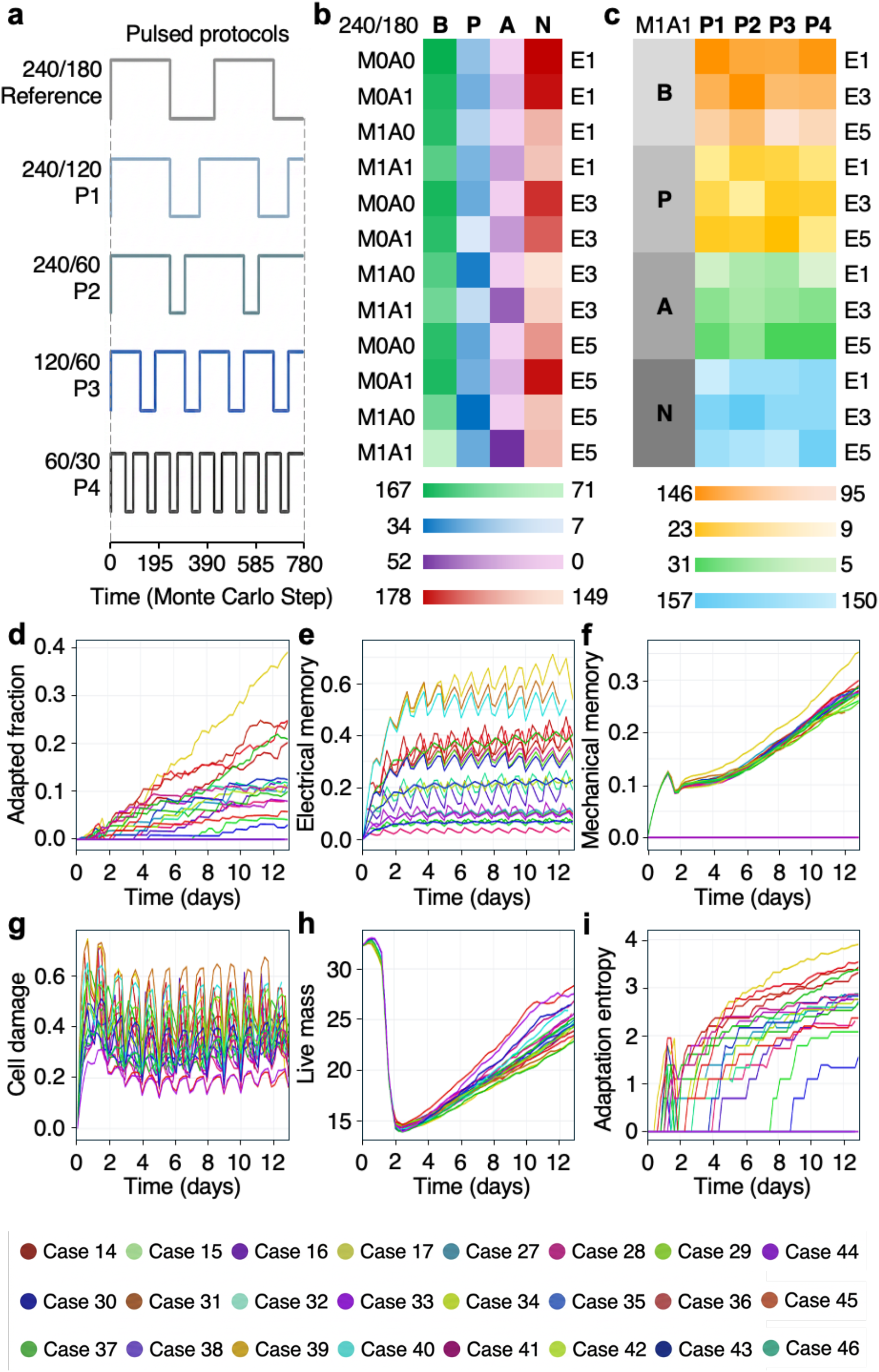
Pulse timing tunes adaptation, memory, and lineage entropy. **(a)** Schematic of the pulsed stimulation protocols; labels denote period/ON duration in Monte Carlo steps: reference 240/180, P1 240/120, P2 240/60, P3 120/60, and P4 60/30. **(b)** Heat map of endpoint state abundance for baseline (B), plastic (P), adapted (A), and necrotic (N) cells across mechanics/adaptation settings and field strengths under the reference protocol. **(c)** Endpoint state abundance for the M1A1 condition across alternative pulse protocols and electric-field amplitudes. **(d-i)** Time courses for the same protocol series showing adapted fraction (**d**), electrical memory (**e**), mechanical memory (**f**), cell damage (**g**), live mass in voxels (**h**), and adaptation entropy (**i**). Pulse structure primarily reshapes adaptation-linked variables while preserving the overall collapse-and-regrowth mass trajectory.

The adaptive channel again exhibited the clearest signature of the intervention. In A0 rows it remained absent, whereas in A1 rows it increased with electric field intensity, with especially pronounced enrichment in M1/A1 at E3 and E5 (Figure 5b). This means that pulsing preserved the same requirement for adaptive competence observed under continuous stimulation but altered the strength of expression of that competence at the endpoint. The protocol-specific comparison within M1/A1 reinforces this point. Across P1–P4, baseline counts ranged from 95 to 146, plastic counts from 9 to 23, adaptive counts from 5 to 31, and necrotic counts remained relatively constrained between 150 and 157 (Figure 5c). In other words, changing duty cycle and switching frequency redistributed the viable phenotypic composition much more than it changed the size of the necrotic compartment. This shows that pulse timing offers an additional control for steering phenotype balance without large changes in gross damage.

The temporal ensemble reveals the mechanisms underlying this broader phenotypic space. Adapted fraction rose from near zero to a broad endpoint distribution of roughly 0.03–0.38, with the upper trajectories separating by about day 4 and continuing upward thereafter (Figure 5d). Electrical memory displayed the strongest oscillatory pattern in the full dataset, climbing through repeated sawtooth cycles to case-specific plateaus between approximately 0.10 and 0.72 (Figure 5e). These oscillations closely mirrored the imposed on/off structure, demonstrating that the memory module remained dynamically entrained to the pulse sequence. Mechanical memory increased more smoothly and accumulated to about 0.25–0.35 in mechanically active cases (Figure 5f), implying that the structural history integrated over longer timescales than electrical exposure. Cell damage oscillated in parallel, spanning on average of 0.15–0.70 (Figure 5g), which indicates repeated cycles of injury and partial repair. Live mass followed the early crash and later regrowth pattern, but endpoint outcomes broadened into an approximately 22–28 range under pulsed forcing (Figure 5h). Finally, adaptation entropy accumulated in staircase-like trajectories from 0 to roughly 1.5–4.0, with the highest values occurring in the same cases that showed the strongest adapted fractions and electrical-memory plateaus (Figure 5i). Pulsed stimulation restructures the timing of memory accumulation and revisits the damage state, thereby supporting a wider spectrum of stable adaptive outcomes compared with continuous electric fields.

## 3. Discussion

This study presents a 3D agent-based framework in which tumor non-genetic diversification, a.k.a. adaptive capacity or phenotypic plasticity, emerges from coupling between vascular oxygenation, electrical stimulation, mechanical interactions, and heritable memory. This study presents a 3D agent-based framework for investigating how coupled physical microenvironmental cues can shape non-genetic phenotypic diversification in a spatially evolving tumor model. By integrating vascular oxygen supply, externally imposed electric field, mechanical crowding and compression cues, stochastic phenotype transitions, and inherited stress memory, the model provides a computational environment in which the consequences of these interacting processes can be examined simultaneously. The emergent quantities of interest are the resulting tissue-level dynamics: spatial phenotype distribution, population composition, growth trajectories, lineage structure, and their dependence on combinations and temporal histories of environmental cues.

In the reference full-model condition, under mid-range applied electric field enabled adaptation (cell transition from a plastic state to a stable adaptive state) and mechanical cues (cell crowding and tissue compression), the tissue followed a reproducible sequence of early damage, necrosis onset, transient live-mass collapse, and partial regrowth with the accumulation of adapted cells.

This pattern supports the idea that short-term stress accommodation can restructure population composition over timescales much shorter than classical clonal evolution, i.e. in the order of seconds to hours, here tested up to 12 days corresponding to ~3000 Monte Carlo Steps (MCS). A key finding is that continuous electrical stimulation mainly altered the balance between growth and adaptation rather than simply increasing endpoint necrosis. Across the continuous-field series, live mass declined and adapted fraction increased significantly with the field amplitude, whereas the final necrotic mass changed comparatively little. In this model, the electrical field therefore behaves less as a purely cytotoxic load and more as a selective and state-redistributing pressure. This distinction important because it suggests that bioelectrical perturbation may alter tumor organization even when gross cell death is not the dominant observable outcome.

The simulations also indicated that mechanics is a major contextual amplifier of the electrical response. When mechanical interactions were disabled, electrical stimulation had a weaker effect on tumor expansion and induced smaller adaptive shifts. When mechanics was enabled, the same electrical forcing produced stronger reductions in live mass and larger increases in adapted state fraction calculated over the total live mass, i.e. alive (non-necrotic) cells in the tumor. This implies that electrical and mechanical stresses jointly shape the effective stress experienced by cells. Adaptive capacity added context by determining whether electrically perturbed cells could transition into a distinct adapted phenotype rather than remaining confined to baseline, plastic, or damaged trajectories. The pulsed simulations extend this interpretation by showing that stimulus timing is itself biologically meaningful in the model. Repeated on/off forcing generated oscillatory electrical memory and damage trajectories, while endpoint composition and adaptation entropy depended on protocol structure. Thus, spheroids in this framework respond not only to the amount of stimulation applied but also to the temporal packaging of that stimulation. This is important for future experimental design because it suggests that amplitude-matched protocols may not be functionally equivalent.

The present model has two main limitations: (i) the electrical, mechanical, and adaptive modules are abstractions of richer biology, and (ii) vascular architecture is fixed. The first point is both a strength and a weakness: on one hand, it allows the isolation of specific cues and quantities regardless of where they originate from (for instance, independently of whether electric field is the result of ionic signaling, cumulative tissue currents or externally applied voltages); on the other hand, it groups diverse electrical contributions into a single end value experienced by cells at the tissue scale. The second point (vasculature and oxygen supply), instead, could be represented as a dynamically remodeled variable as a function of cell states in future studies to emulate angiogenesis. Despite these limitations, the model is useful for deployment in hypothesis-generation/testing to enrich the design of experimental studies where coupled microenvironmental pressures produce emergent spatial heterogeneity. It identifies electric fields and mechanics as control variables for future investigations on cellular adaptation in cancer. More broadly, cellular adaptation dynamics interlinked with electrical and mechanical cues are observed in multiple other processes, including aging, autoimmune disorders, development, morphogenesis, and fertility.

## 4. Methods

### Computational framework

Tumor evolution was simulated in CompuCell3D v4.7.0 on a 150×150×50 lattice with no-flux boundaries in all spatial directions for 3000 Monte Carlo steps (MCS). The CellType plugin defined medium M and four cellular states: baseline tumor cells B, plastic tumor cells P, adapted tumor cells A, and necrotic cells N. The Cellular Potts temperature was T_CPM_=10, and the contact-neighbor order was 2. One MCS was mapped to 6 min for reporting, in accordance with previous studies^36^, and one lattice voxel represented 10 μm per side. The implementation was organized into steppables for tissue initialization, vascular/electrical field updates, mechanical cue evaluation, phenotype and fate updates, mitosis, and output generation. Within each MCS, field updates were applied first, followed by mechanical cue computation, phenotype/damage/memory updates, mitosis, and output writing. Cell dynamics followed a 3D Cellular Potts formulation in which copy attempts were accepted with probability

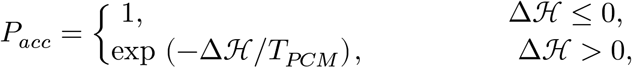

The Hamiltonian was

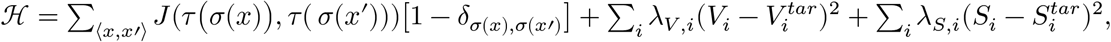

where *V*_*i*_ and *S*_*i*_ are cell volume and surface, 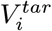 and 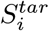 are their targets, and *J* is the contact-energy matrix. Using the ordering (M, B, P, A, N), the implemented contact energies are

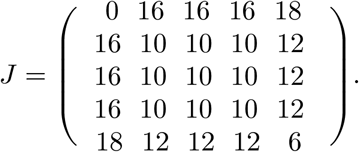

Live cells used λ_*V*_ = 35, λ_*S*_ = 3, and *S*^*tar*^ = 150, whereas necrotic cells used λ_*V*_ = 80, λ_*S*_ = 4, and *S*^*tar*^ = 90. Cell target volume was initialized at 125 voxels.

### Tumor initialization and inherited heterogeneity

The initial tumor was generated near the center of the domain as a compact aggregate of baseline cells. Specifically, 5×5×5-voxel B cells were packed within a spherical region of radius 20 voxels centered at the lattice midpoint, with placements excluded if they overlapped vessel voxels. Each initial cell was assigned one of 16 founder identifiers according to its azimuthal angle around the aggregate center in the x-y plane. All initial cells began with zero electrical, mechanical, and hypoxic memory, zero accumulated damage, zero hypoxia clock, no prior phenotype switching, and no adaptation lineage assignment. To introduce inheritable non-genetic heterogeneity, each initial cell was assigned an electrical sensitivity, mechanical sensitivity, plasticity bias, repair bias, growth bias, and lock-in bias. These were sampled independently from clipped Gaussian distributions centered at 1.0, with standard deviations 0.10, 0.10, 0.15, 0.10, 0.10, and 0.10, respectively. Clipping bounds were [0.5,1.5] for all traits except for the growth bias, which used [0.6,1.4]. Cell-specific division thresholds were initialized as 200η_i_, where η_i_~N(1, 0.05^2^) clipped to [0.85,1.15]. Initial birth times were staggered over the preceding 140 MCS to desynchronize early divisions.

### Oxygen field

Oxygen was volved by diffusion-decay using DiffusionSolverFE in CompuCell3D, defined as

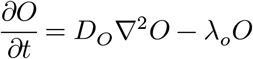

with *D*_*O*_ = 0.15 and λ_*o*_ = 10^−4^, under no-flux boundary conditions. Vessel source voxels were defined as z-spanning cylindrical columns of radius 2 voxels placed on a jittered square grid in the x-y plane with spacing 45 voxels, outer margin 25 voxels, and random jitter ±5 voxels. The initial background oxygen level was 0.25, and vessel voxels were reset to O=1.0 after each update. After each field update, oxygen was decreased on cell-occupied voxels by a phenotype-specific uptake term:

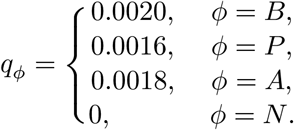

Cell-level oxygen *O*_*i*_(*t*) was sampled at the cell center of mass. The hypoxic stress signal used in damage, memory, and switching was

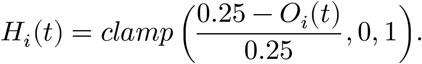

### Electrical stimulation

Unlike oxygen, voltage was stored in a non-diffusing scalar field and overwritten directly each MCS rather than being evolved by diffusion-decay. In the XML configuration, the voltage field had diffusion and decay constants set to zero. Let *I*_*E*_(*t*) denote the stimulation indicator:

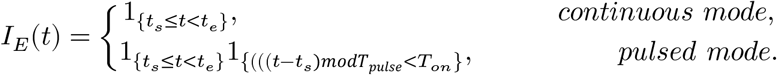

During on intervals, the extracellular voltage field was imposed as a left-to-right linear profile:

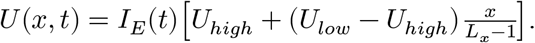

Left and right electrode regions were implemented as 8-voxel-wide lateral strips, offset by 12 voxels from the x boundaries and spanning the y direction except for 12-voxel margins, and were clamped to *U*_*high*_ and *U*_*low*_, respectively. During off intervals, *U*(*x, t*) = 0. The sampled local voltage *U*_*i*_(*t*) was stored as an output, but the phenotype rules used the field-gradient magnitude. Because the imposed profile is linear, the raw electrical cue was spatially uniform during on phases:

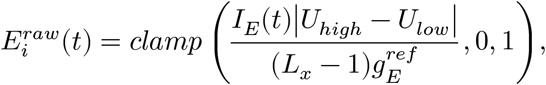

where 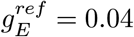 Cell-specific electrical exposure was then

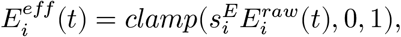

where 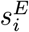 is the inherited electrical sensitivity of cell *i*.

### Mechanical cue

Mechanical cueing was computed for each live cell from a weighted combination of compression and crowding. When the cue was enabled, a cell compression component was

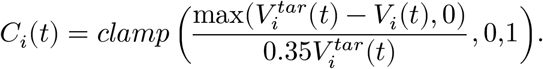

Let 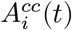 and 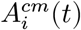 be the cell-cell and cell-medium contact areas, respectively. The crowding fraction was

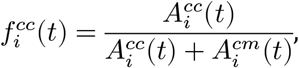

and the normalized crowding term was

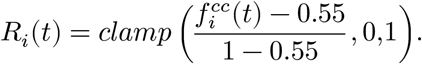

The raw mechanical cue was then

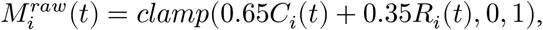

and effective mechanical cue used by phenotype rules was

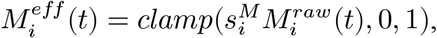

where 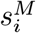 is the inherited mechanical sensitivity. When the mechanical module was disabled,

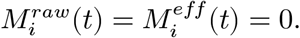

*Damage, stress memory, and hypoxia*. The oxygen-support factor used in growth was

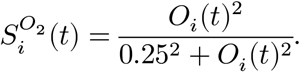

Damage was updated before phenotype switching. For phenotype *ϕ* ∈ {*B, P*, *A*},

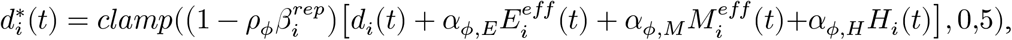

where 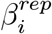 is a repair bias, and

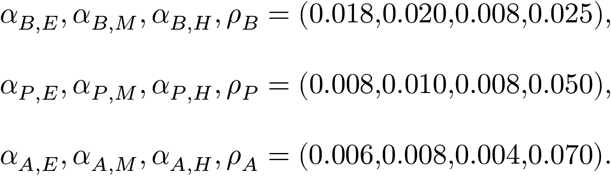

Electrical, mechanical, and hypoxic memory components were updated as

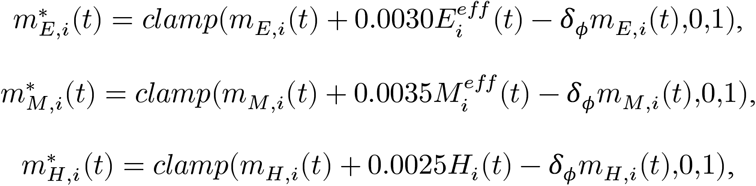

With δ_*B*_ = 0.0040, δ_*P*_ = 0.0015, and δ_*A*_ = 0.0018. Total memory was then:

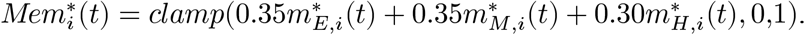

A hypoxia clock was also maintained:

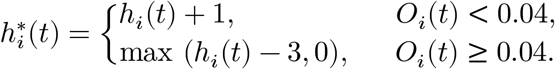

If 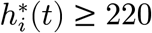, the cell was converted to the necrotic state before phenotype switching.

### Phenotype switching

Phenotype transitions were modeled as stochastic Bernoulli events evaluated once per MCS for surviving live cells. For baseline cells,

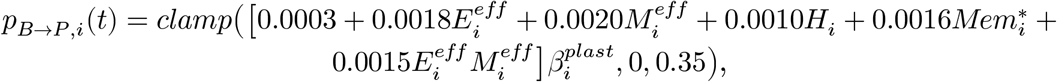

where 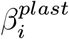 is a plasticity bias. If the transition occurred, the cell entered the plastic state and its plastic-state residence clock was initialized to 1. For plastic cells, the plastic-state residence clock was incremented each MCS and reversion to baseline was tested first. Defining a stress-relief factor as:

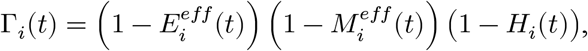

the reversion probability was:

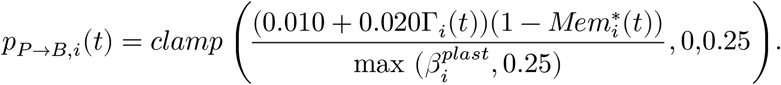

If a *P* → *B* transition did not occur, *P* → *A* was then tested. Let τ_i_^*P*^ (*t*) denote the plastic-state residence clock and define:

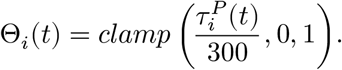

The adaptation probability was:

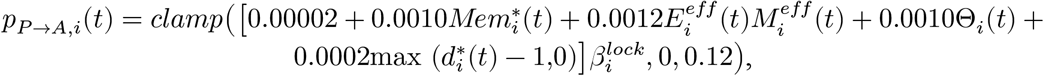

where 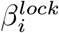 is a lock-in bias. When a cell first entered the adapted state, it was assigned a new adaptation-lineage identifier that was inherited by its descendants. For adapted cells, the reverse transition was

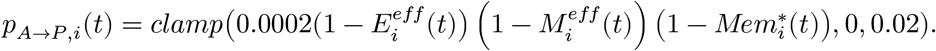

### Growth, mitosis, and cell death

After any phenotype change within the same MCS, growth was assigned according to the resulting live phenotype. For B, P, and A cells, respectively,

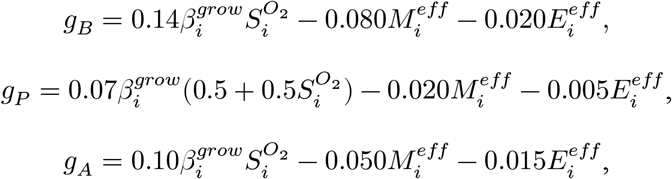

where 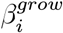 is a growth bias. An additional damage penalty was applied when 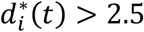,

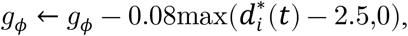

and an additional hypoxia penalty of −0.12 was applied when *O*_*i*_ (*t*) < 0.04. Target volume was then updated as

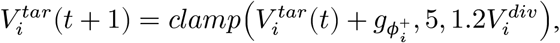

where 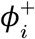 is the post-switch phenotype and 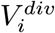is the cell-specific division threshold. Damage-dependent death probabilities were:

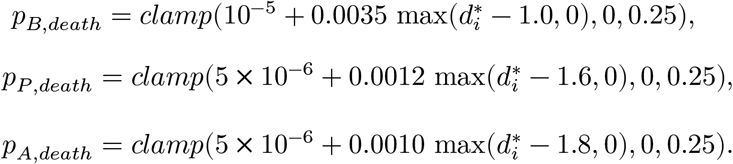

If death occurred, the cell was converted to the necrotic state. At the moment of death, the necrotic target volume was set to max(4,0.6V_i_), with λ_v_=80, λ_s_=4, and 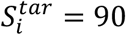. Necrotic target volume was then reduced by 0.10 per MCS, and necrotic cells were deleted once their actual volume reached ≤ 3 voxels. A live cell divided when 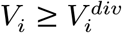 andits aged satisfied a phenotype-specific minimum cycle time: *a*_*min,B*_ = 140, *a*_*min,P*_ = 320, *a*_*min,A*_ = 190 MCS. Division was performed with random orientation. At mitosis, both post-mitotic cells retained the parental phenotype, founder identity, and adaptation identity, and the child recorded the parent cell ID. Stress memory and damage were inherited as:

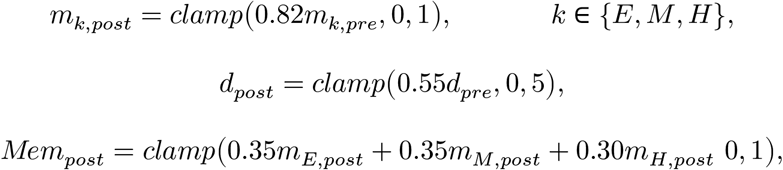

and the hypoxia clock was halved:

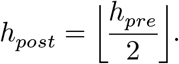

Parent and child target volumes were reset to 125. The generation number and cumulative division count stored in each cell were both incremented by one relative to the pre-mitotic cell. Cell-specific division thresholds were inherited with bounded multiplicative Gaussian variation around the parental value (standard deviation 0.03, multiplicative clipping [0.9,1.1]). Electrical sensitivity, mechanical sensitivity, plasticity bias, repair bias, growth bias, and lock-in bias were inherited with small multiplicative log-normal jitter (Gaussian log-space standard deviation 0.02) and then clipped to their allowed ranges.

### Simulation outputs and readouts

Summary tables were written every 50 MCS. Full cell-level and field-level snapshots were exported every 2000 MCS. Orthogonal mid-slice images and 3D voxel renderings of the cell field, cell volume, oxygen, and voltage were exported every 100 MCS, and diagnostic plots were refreshed every 500 MCS. Cell-level outputs included phenotype, lineage identifiers, age, memory, damage, local oxygen and voltage, raw and effective electrical and mechanical cues, sensitivities, division thresholds, volume, target volume, and center-of-mass coordinates. Aggregate outputs included phenotype counts and fractions, occupied live and necrotic volumes, constant-density mass proxies, mean cue values, mean oxygen and voltage, founder and adaptation-lineage Shannon entropies, number of adaptation lineages, projected areas, geometric spans, a 90th-percentile live-cell radius, live center of mass, and cumulative event counters for divisions, phenotype transitions, and death modes.

The readouts were defined as follows. Let

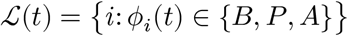

denote live cells, and

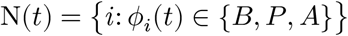

denote necrotic cells. Live, necrotic, and total mass were reported as constant-density mass proxies proportional to occupied volume:

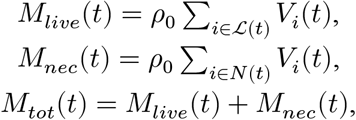

with constant *ρ*_0_. State fractions *f*_*B*_(*t*), *f*_*P*_ (*t*), *f*_*A*_(*t*) were computed over live cells only. The adapted fraction was

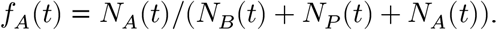

The hypoxic fraction was

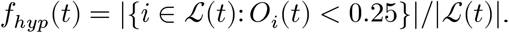

The q90 radius was the 90th percentile of live-cell distances from the live-cell center of mass. Adaptation entropy was the Shannon entropy of adaptation-lineage identifiers among adapted cells,

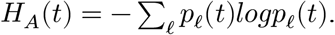

Unless otherwise stated, plotted memory components, integrated memory, and cell damage were live-cell means. Volumes were converted using 1 voxel = 10^3^ μm^3^. Landmark times were defined as T1, the first MCS with nonzero necrotic mass; T2, the MCS of minimum live mass after T1; and T3, the final simulated MCS.

### Sample case

The above rules were applied to all simulated cases, presented in **Table 1**. Here, Case 34 is taken as a sample case. The random seed was 41, and both Python and NumPy random number generators were initialized with this seed. Electrical stimulation and mechanical cueing were enabled, and adaptation was enabled by retaining nonzero P→A transition coefficients. The electric field operated in pulsed mode with *t*_*s*_ = 0, *t*_*e*_ = 3000,

**Table 1.**
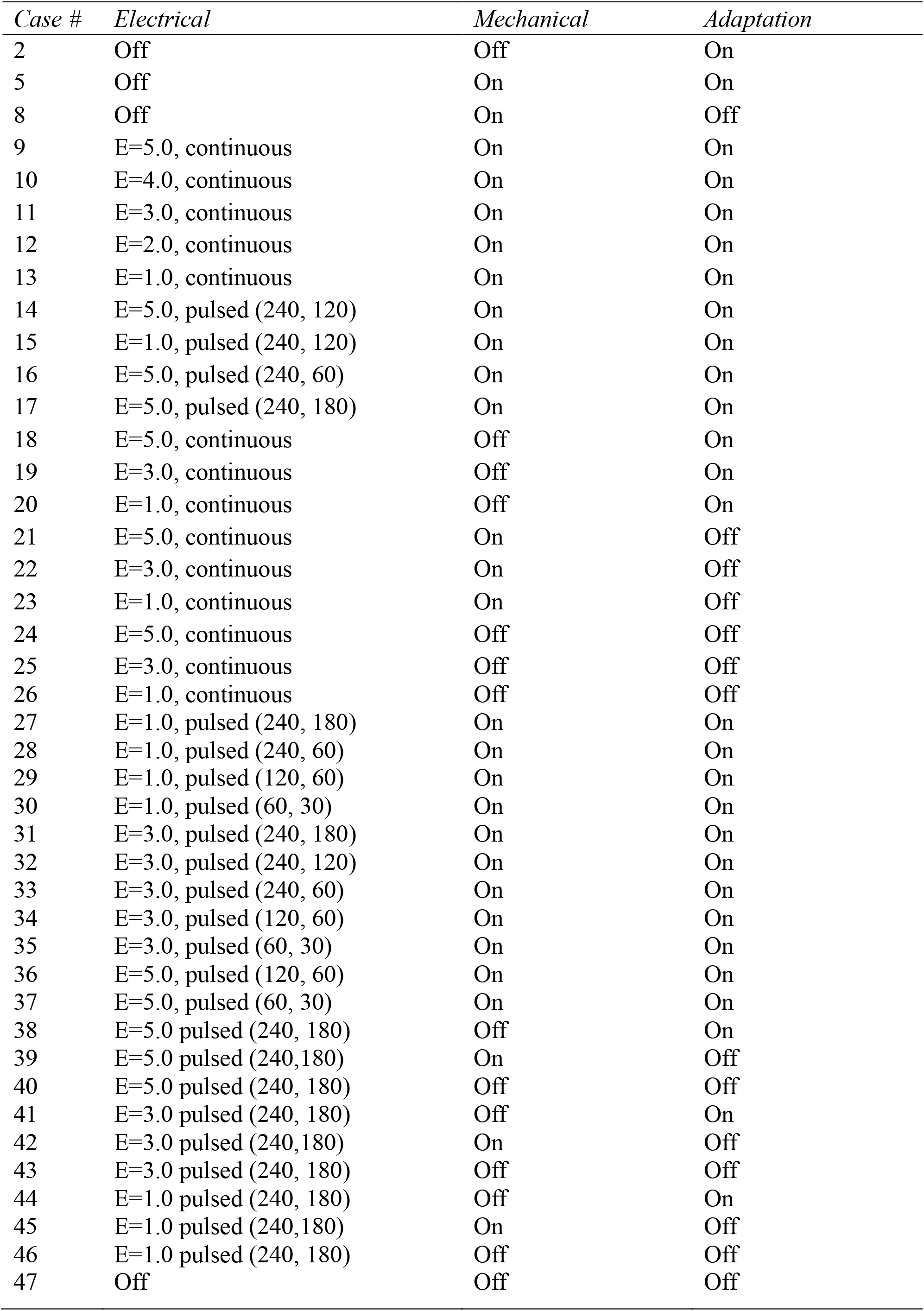
List of the simulation cases presented in this work.

| *Case #* | *Electrical* | *Mechanical* | *Adaptation* |
| --- | --- | --- | --- |
| 2 | Off | Off | On |
| 5 | Off | On | On |
| 8 | Off | On | Off |
| 9 | E=5.0, continuous | On | On |
| 10 | E=4.0, continuous | On | On |
| 11 | E=3.0, continuous | On | On |
| 12 | E=2.0, continuous | On | On |
| 13 | E=1.0, continuous | On | On |
| 14 | E=5.0, pulsed (240, 120) | On | On |
| 15 | E=1.0, pulsed (240, 120) | On | On |
| 16 | E=5.0, pulsed (240, 60) | On | On |
| 17 | E=5.0, pulsed (240, 180) | On | On |
| 18 | E=5.0, continuous | Off | On |
| 19 | E=3.0, continuous | Off | On |
| 20 | E=1.0, continuous | Off | On |
| 21 | E=5.0, continuous | On | Off |
| 22 | E=3.0, continuous | On | Off |
| 23 | E=1.0, continuous | On | Off |
| 24 | E=5.0, continuous | Off | Off |
| 25 | E=3.0, continuous | Off | Off |
| 26 | E=1.0, continuous | Off | Off |
| 27 | E=1.0, pulsed (240, 180) | On | On |
| 28 | E=1.0, pulsed (240, 60) | On | On |
| 29 | E=1.0, pulsed (120, 60) | On | On |
| 30 | E=1.0, pulsed (60, 30) | On | On |
| 31 | E=3.0, pulsed (240, 180) | On | On |
| 32 | E=3.0, pulsed (240, 120) | On | On |
| 33 | E=3.0, pulsed (240, 60) | On | On |
| 34 | E=3.0, pulsed (120, 60) | On | On |
| 35 | E=3.0, pulsed (60, 30) | On | On |
| 36 | E=5.0, pulsed (120, 60) | On | On |
| 37 | E=5.0, pulsed (60, 30) | On | On |
| 38 | E=5.0 pulsed (240, 180) | Off | On |
| 39 | E=5.0 pulsed (240,180) | On | Off |
| 40 | E=5.0 pulsed (240, 180) | Off | Off |
| 41 | E=3.0 pulsed (240, 180) | Off | On |
| 42 | E=3.0 pulsed (240,180) | On | Off |
| 43 | E=3.0 pulsed (240, 180) | Off | Off |
| 44 | E=1.0 pulsed (240, 180) | Off | On |
| 45 | E=1.0 pulsed (240,180) | On | Off |
| 46 | E=1.0 pulsed (240, 180) | Off | Off |
| 47 | Off | Off | Off |

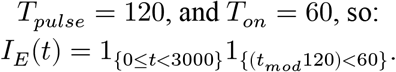

The applied voltages were *U*_*high*_ = 3.0 and *U*_*low*_ = 0.0, with 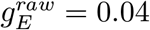. Because *L*_*x*_ = 150, the raw electrical cue during on pulses, before scaling by the cell-specific electrical sensitivity, was:

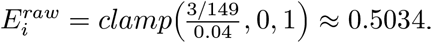

All other cases retain the same equations and update order, differing only by parameter toggles controlling electrical stimulation, mechanical cueing, adaptation probabilities, pulse schedule, and field amplitude.

## 5. Conclusion

This work establishes a modular 3D agent-based framework for studying how oxygen availability, mechanical context, electric fields, and inherited stress history interact to shape tumor population dynamics. The simulations predict that the consequences of electric fields depend on the surrounding mechanical and vascular context, and that different temporal stimulation protocols can produce distinct trajectories of phenotypic composition and electrical/mechanical memory. These predictions provide experimentally testable hypotheses and a computational foundation for future incorporation of calibrated electrophysiological, mechanical, and molecular mechanisms. More broadly, the study offers a preliminary demonstration that physical oncology should treat stress as a dynamic selection environment that can redirect tumour evolution rather than only suppress growth. Because the model distinguishes reversible perturbation from durable ecological change, it provides a basis for testing interventions designed to prevent stable adaptation, exploit sensitive states, or uncouple viability from persistence.

## Data availability statement

The codes from which all data was generated are available on GitHub: https://github.com/moreddurosalia-code/Electromechanical-Adaptation-Modeling

## Notes

https://github.com/moreddurosalia-code/Electromechanical-Adaptation-Modeling

### Competing Interest Statement

The authors have declared no competing interest.

## References

1. de Visser, K. E.; Joyce, J. A. The evolving tumor microenvironment: From cancer initiation to metastatic outgrowth. Cancer Cell 2023, 41 (3), 374–403. DOI: 10.1016/j.ccell.2023.02.016.

2. Khelifa, L.; Hu, Y.; Tall, J.; Khelifa, R.; Ali, A.; Poon, E.; Khelifa, M. Z.; Yang, G.; Jones, C.; Moreddu, R.,,,,, et al. Diagnostic technologies for neuroblastoma. Lab Chip 2025, 25 (15), 3630–3664. DOI: 10.1039/d4lc00005f.

3. Marusyk, A.; Polyak, K. Tumor heterogeneity: causes and consequences. Biochim Biophys Acta 2010, 1805 (1), 105–117. DOI: 10.1016/j.bbcan.2009.11.002.

4. Franco Jones, C.; Dias, D.; Moreira, A. C.; Goncalves, G.; Cinti, S.; Djamgoz, M. B. A.; Castelo Ferreira, F.; Sanjuan-Alberte, P.; Moreddu, R. Multilevel classification framework for breast cancer cell selection and its integration with advanced disease models. iScience 2025, 28 (10), 113579. DOI: 10.1016/j.isci.2025.113579.

5. Senft, D.; Ronai, Z. e. A. Adaptive Stress Responses During Tumor Metastasis and Dormancy. Trends in Cancer 2016, 2 (8), 429–442. DOI: 10.1016/j.trecan.2016.06.004.

6. Franca, G. S.; Baron, M.; King, B. R.; Bossowski, J. P.; Bjornberg, A.; Pour, M.; Rao, A.; Patel, A. S.; Misirlioglu, S.; Barkley, D.,,,,, et al. Cellular adaptation to cancer therapy along a resistance continuum. Nature 2024, 631 (8022), 876–883. DOI: 10.1038/s41586-024-07690-9.

7. Hanahan, D. Hallmarks of cancer-Then and now, and beyond. Cell 2026, 189 (8), 2254–2277. DOI: 10.1016/j.cell.2025.12.049.

8. Kulkarni, P.; Ramisetty, S.; Bruno, D.; Tan, T.; Merla, A.; Salgia, R. Phenotypic Plasticity, Non-genetic Mechanisms, and Immune Drug Resistance in Cancer. Cancer Treat Res 2025, 129, 309–324. DOI: 10.1007/978-3-031-97242-3_14.

9. Cheung, K. J.; Horne-Badovinac, S. Collective migration modes in development, tissue repair and cancer. Nat Rev Mol Cell Biol 2025, 26 (10), 741–758. DOI: 10.1038/s41580-025-00858-9.

10. Li, J.; Ravindran, P. T.; O’Farrell, A.; Busch, G. T.; Boe, R. H.; Niu, Z.; Woo, S.; Dunagin, M. C.; Jain, N.; Goyal, Y.,,,,, et al. AP-1 mediates cellular adaptation and memory formation. Nat Commun 2026, 17 (1). DOI: 10.1038/s41467-026-70862-w.

11. Franca, G. S.; Yanai, I. A mechanism for adaptive genome regulation in cancer. Nature 2026, 652 (8110), 581–590. DOI: 10.1038/s41586-026-10269-1.

12. Moreddu, R. Bioinspired Engineering beyond Homeostasis. Advanced Intelligent Systems 2025, 8 (1). DOI: 10.1002/aisy.202500435.

13. Greulich, P.; Levin, M.; Moreddu, R. Oncomorphic neural agent populations for resource-limited sequential learning. arXiv e-prints 2025, arXiv: 2503.12743.

14. Chen, Z.; Han, F.; Du, Y.; Shi, H.; Zhou, W. Hypoxic microenvironment in cancer: molecular mechanisms and therapeutic interventions. Signal Transduct Target Ther 2023, 8 (1), 70. DOI: 10.1038/s41392-023-01332-8.

15. Bloch, N.; Harel, D. The tumor as an organ: comprehensive spatial and temporal modeling of the tumor and its microenvironment. BMC Bioinformatics 2016, 17 (1), 317. DOI: 10.1186/s12859-016-1168-5.

16. Zhu, Y.; Chen, J.; Chen, C.; Tang, R.; Xu, J.; Shi, S.; Yu, X. Deciphering mechanical cues in the microenvironment: from non-malignant settings to tumor progression. Biomark Res 2025, 13 (1), 11. DOI: 10.1186/s40364-025-00727-9.

17. Moreddu, R.; Mahmoodi, N.; Kassanos, P.; Vigolo, D.; Mendes, P. M.; Yetisen, A. K. Stretchable Nanostructures as Optomechanical Strain Sensors for Ophthalmic Applications. ACS Applied Polymer Materials 2021, 3 (11), 5416–5424. DOI: 10.1021/acsapm.1c00703.

18. Boschi, A.; Iachetta, G.; Buonocore, S.; Hubarevich, A.; Moreddu, R.; Dipalo, M.,,,,, et al. Interferometric Biosensor for High Sensitive Label-Free Recording of HiPS Cardiomyocytes Contraction in Vitro. Nano Lett 2024, 24 (22), 6451–6458. DOI: 10.1021/acs.nanolett.3c04291.

19. Tan, M.; Song, B.; Zhao, X.; Du, J. The role and mechanism of compressive stress in tumor. Front Oncol 2024, 14, 1459313. DOI: 10.3389/fonc.2024.1459313.

20. Kalukula, Y.; Luciano, M.; Simanov, G.; Charras, G.; Brückner, D. B.; Gabriele, S. The actin cortex acts as a mechanical memory of morphology in confined migrating cells. Nature Physics 2025, 21 (9), 1451–1461. DOI: 10.1038/s41567-025-02980-z.

21. Stylianopoulos, T.; Martin, J. D.; Snuderl, M.; Mpekris, F.; Jain, S. R.; Jain, R. K. Coevolution of solid stress and interstitial fluid pressure in tumors during progression: implications for vascular collapse. Cancer Res 2013, 73 (13), 3833–3841. DOI: 10.1158/0008-5472.CAN-12-4521.

22. Rizzo, M. G.; Fazio, E.; De Pasquale, C.; Sciuto, E. L.; Cannata, G.; Multisanti, C. R.; Impellitteri, F.; D’Agostino, F. G.; Guglielmino, S. P. P.; Faggio, C.,,,,, et al. Physiopathological Features in a Three-Dimensional In Vitro Model of Hepatocellular Carcinoma: Hypoxia-Driven Oxidative Stress and ECM Remodeling. Cancers (Basel) 2025, 17 (18). DOI: 10.3390/cancers17183082.

23. Cadinu, P.; Burgess, M. K.; Franco Jones, C.; Iarossi, M.; Schröter, M.; Nakatsuka, N.; Djamgoz, M. B. A.; Gonçalves, G.; Abayzeed, S.; Sanjuán-Alberte, P.,,,,, et al. Bioelectrical Interfaces Beyond Excitable Cells: Cancer, Aging, and Gene Expression Modulation. Advanced Materials Interfaces 2026, 13 (10). DOI: 10.1002/admi.202500999.

24. Melikov, R.; De Angelis, F.; Moreddu, R. High-frequency extracellular spiking in electrically-active cancer cells. bioRxiv 2024, 2024.2003.2016.585162. DOI: 10.1101/2024.03.16.585162.

25. Moreddu, R. Nanotechnology and Cancer Bioelectricity: Bridging the Gap Between Biology and Translational Medicine. Adv Sci (Weinh) 2024, 11 (1), e2304110. DOI: 10.1002/advs.202304110.

26. Kyndiah, A.; Zemignani, G. Z.; Ronchi, C.; Tullii, G.; Khudiakov, A.; Iachetta, G.; Chiodini, S.; Moreddu, R.; Viola, F. A.; Schwartz, P. J.,,,,, et al. Non-invasive action potential recordings using printed electrolyte-gated polymer field-effect transistors. Nat Commun 2025, 16 (1), 8143. DOI: 10.1038/s41467-025-63484-1.

27. Abayzeed, S.; Galvis, D.; Medel, K. R.; Jones, C. F.; Gonzalez, O. B.; Setchfield, K.; Moreddu, R.; Somekh, M. G.; Wedgwood, K. C. A.; Smith, P. Plasmonic imaging of living pancreatic beta-cell networks. Sci Rep 2026, 16 (1), 3993. DOI: 10.1038/s41598-025-34094-0.

28. Moreddu, R.; Boschi, A.; d’Amora, M.; Hubarevich, A.; Dipalo, M.; De Angelis, F. Passive Recording of Bioelectrical Signals from Non-Excitable Cells by Fluorescent Mirroring. Nano Lett 2023, 23 (8), 3217–3223. DOI: 10.1021/acs.nanolett.2c05053.

29. Bearer, E. L.; Lowengrub, J. S.; Frieboes, H. B.; Chuang, Y. L.; Jin, F.; Wise, S. M.; Ferrari, M.; Agus, D. B.; Cristini, V. Multiparameter computational modeling of tumor invasion. Cancer Res 2009, 69 (10), 4493–4501. DOI: 10.1158/0008-5472.CAN-08-3834.

30. Deisboeck, T. S.; Wang, Z.; Macklin, P.; Cristini, V. Multiscale cancer modeling. Annu Rev Biomed Eng 2011, 13, 127–155. DOI: 10.1146/annurev-bioeng-071910-124729.

31. Simpson, M. J.; Buenzli, P. R. Dimension-dependent continuum limits in tissue mechanics. arXiv preprint arXiv:2607.07000 2026.

32. Morris, M. K.; Saez-Rodriguez, J.; Sorger, P. K.; Lauffenburger, D. A. Logic-based models for the analysis of cell signaling networks. Biochemistry 2010, 49 (15), 3216–3224. DOI: 10.1021/bi902202q.

33. Wang, Y.; Casarin, S.; Daher, M.; Mohanty, V.; Dede, M.; Shanley, M.; Dondossola, E.; La Posta, L.; Basar, R.; Rezvani, K.,,,,, et al. Agent-based modeling of cellular dynamics in adoptive cell therapy. Commun Biol 2026, 9 (1). DOI: 10.1038/s42003-026-09653-4.

34. van Genderen, M. N. G.; Kneppers, J.; Zaalberg, A.; Bekers, E. M.; Bergman, A. M.; Zwart, W.; Eduati, F. Agent-based modeling of the prostate tumor microenvironment uncovers spatial tumor growth constraints and immunomodulatory properties. NPJ Syst Biol Appl 2024, 10 (1), 20. DOI: 10.1038/s41540-024-00344-6.

35. Baselga-Cervera, B.; Medina-Chavez, N. O.; Gettle, N.; Travisano, M. Stochastic phenotypic switching arises in response to directional selection in experimentally evolved multicellular yeast. Commun Biol 2025, 9 (1), 134. DOI: 10.1038/s42003-025-09414-9.

36. Swat, M.H.; Thomas, G.L.; Shirinifard, A.; Clendenon, S.G.; Glazier, J.A. Emergent Stratification in Solid Tumors Selects for Reduced Cohesion of Tumor Cells: A Multi-Cell, Virtual-Tissue Model of Tumor Evolution Using CompuCell3D. PLoS One 2015, 10 (6), e0127972. DOI: 10.1371/journal.pone.0127972.

